# Hexadecimal data encryption in paranemic crossover (PX) DNA

**DOI:** 10.64898/2026.07.17.738806

**Authors:** Anna Karpen, Arun Richard Chandrasekaran

## Abstract

DNA is highly programmable and efficient for encoding information. In this work, we use the paranemic crossover (PX) DNA structures for a binary-encoded system. As substrates for data storage, we designed a combination of PX and anti-PX structures where the four strands of the PX motif are complementary to those in the anti-PX motif. To write data, we programmed encoding elements in each of the four strands of the PX and anti-PX motifs. The encoded data remains encrypted until the samples are processed at a specific temperature, when the PX and anti-PX motifs reassociate to four distinct duplexes, defined by the encoding elements and retrieved using an electrophoretic readout. We show that the encoded information is stable for several days when stored at 20 °C, 37 °C or outdoors, with the encrypted structures showing higher nuclease resistance compared to the decrypted structures. Using this strategy, we demonstrate hexadecimal encoding using a combination of 4 bits, encrypting specific words and color codes. We envision such systems could find use barcoding, secure messaging and authentication.

---

In the 2015 Hollywood movie *The Martian*, Matt Damon’s astronaut-scientist character Mark Watney is stranded on Mars. Limited by the ability to communicate with his colleagues at NASA, he has a sudden idea and exclaims “*Hexadecimals to the rescue*”. He then uses the hexadecimal coding system to communicate with the space station a series of words and sentences. Such binary codes are prevalent in computers and instruments in everyday usage. Recently, binary encoding has also been expanded to DNA-based systems where data can be stored, processed, transmitted and read out. ^1^ The sequence programmability and long term stability of DNA are key features that make DNA useful in data storage^2–4^ and molecular computation.^5^ DNA has been used for archival storage of information such as books,^6^ images,^6^ and Shakespearean sonnets.^7^ Large-scale synthesis of DNA at low cost, high yields, and at longer lengths has contributed to these developments.^8^ Data stored or encrypted in DNA sequences typically use DNA sequencing to read out the encoded information, and more recently through nanopore technologies capable of direct single-molecule readouts.^9^ Molecular encryption in such DNA storage systems has also been achieved by chemical functionalization and biomolecular processes such as the incorporation of 2-aminoadenine, an adenosine analog that cannot be faithfully replicated by DNA polymerase,^10^ embedding a chimeric D-DNA/L-DNA key molecule in a D-DNA storage library,^11^ and by epigenetic modifications on pre-made DNA templates.^12,13^

While typical DNA-based data storage methods encode information in sequences of DNA strands, the structures of DNA can also be used to encode information for short term storage^14^ or encryption.^15^ These encoding systems have used simple structures such as single stranded or double stranded DNA^16^ and hairpins,^17^ and more complex structures such as multi-crossover DNA motifs,^18^ DNA origami^15,19^ and core-shell DNA condensates.^20^ Typically data encoding and processing involves the binding of additional biomolecules such as DNA,^21^ RNA,^22^ or enzymes^23^ as inputs on existing DNA nanostructures. The presence or absence of the inputs define the encoded information, and a combination of such inputs provides multi-bit informational storage. In some cases, the biomolecular inputs cause a structural reconfiguration that provides different structural conformations as outputs that translate to bits.^24^ Our own prior work in this area has used a multi-state DNA nanoswitch that can encode bits as looped states when bound to specific DNA or RNA inputs.^24–26^ Further, such information storage systems allow data encryption, where the stored information is read out correctly only in the presence of a decryption key, with some examples of decryption keys including DNA or RNA (via strand displacement),^24^ enzymes (via cleavage or extension)^22,27^ and light (via cleavage or dehybridization).^27,28^ The encrypted information can be read out through gel imaging,^25^ atomic force microscopy,^15^ fluorescence changes,^21^ DNA-PAINT analysis^29^ or confocal microscopy.^20^ Here we present a new strategy for DNA data encoding using the paranemic crossover (PX) DNA structure. We demonstrate encryption of data within the component strands of PX DNA and decryption by temperature-dependent reassociation of PX DNA into size-defined duplexes that provide 4 bits, allowing storage and relay of information represented in hexadecimal codes.

The PX DNA structure contains four strands hybridized into two double helical domains that are connected by strand crossovers at every possible point where the two helices come into proximity (**Fig. 1a, Fig. S1** and **Table S1**). PX DNA motifs have been used in the construction of polyhedral DNA objects,^30^ 1-dimensional and 2-dimensional arrays^31,32^ and nanodevices with rotatable helical domains.^33,34^ We, and others, have also developed reconfigurable PX devices that can respond to DNA, RNA, or enzymes.^35–38^ In our recent work,^39^ we showed that PX and anti-PX structures can reassociate at specific temperatures into duplexes. This concept provides two features that are useful for molecular encryption. First, it allows information encoding in PX structures using 4 unique strands, in combination with another 4 strands in the anti-PX structure. Second, the information encoded in the PX structure remains encrypted, as any changes to the structure is triggered only at a specific temperature. In this work, we use this model system to encrypt information as 4-bit binary codes that allow hexadecimal data storage, decrypt by treating the encoded structures at a specific temperature, and readout the decrypted multi-bit information using an electrophoretic readout.

**Figure 1.**
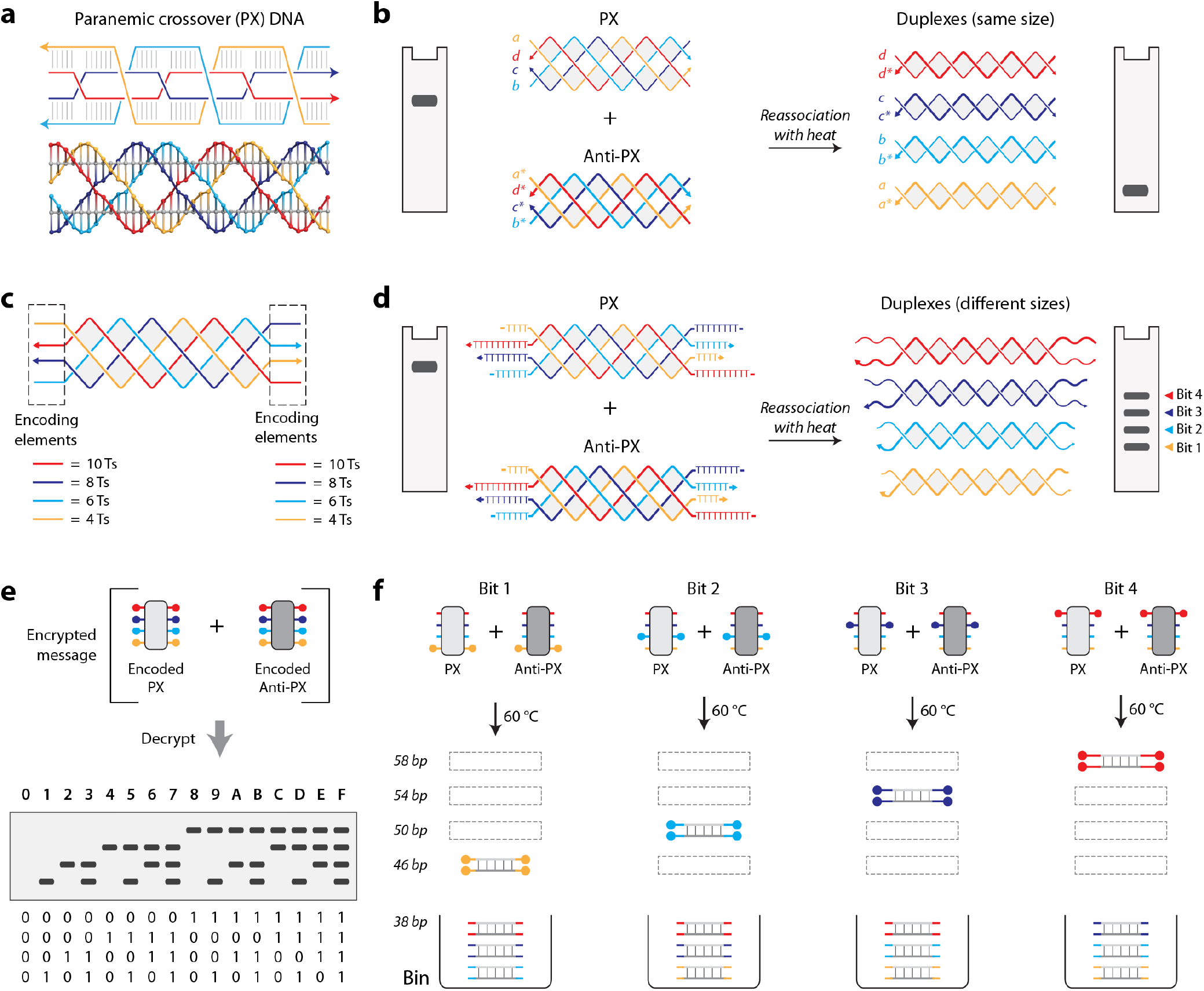
Data encryption strategy using paranemic crossover (PX) DNA. (a) Schematic and molecular model of PX DNA. (b) Reassociation of PX DNA into duplexes when treated with heat. (c) PX DNA with encoding elements in each of the four strands. Each strand contained additional 4, 6, 8 or 10 Ts on either terminus. (d) Reassociation of PX DNA with encoding elements into four distinct duplexes, allowing four encoded bits. (e) PX encryption for hexadecimal readout. (f) Illustration showing the encoding of individual bits using PX DNA, decryption using heat, and the obtained products.

The PX and anti-PX motifs we had previously used reassociate at a specific temperature to form four duplexes (**Fig. 1b**). The four component strands of the anti-PX structure are complementary to the four strands in the PX molecule. That is, each strand in the PX (a, b, c, and d) has a fully complementary partner in the anti-PX (a*, b*, c* and d*, respectively). When mixed and heated, the PX and anti-PX motifs result in the formation of four duplexes of the same molecule weight (38 bp each), thus resulting in a single band on the gel after reassociation (**Fig. 1b**). We modified the termini of the four strands of the PX and anti-PX molecules to add “encoding elements” that can distinguish the product duplexes (**Fig. 1c**). To obtain a 4-bit readout (i.e. 4 unique bands addressable by specific encoding elements), we redesigned the PX and anti-PX strands to be composed of strands that contain additional thymines at the termini (**Fig. 1d**). To obtain four distinct outputs, we added a different number of Ts to each terminus of the strands in the PX and anti-PX: 4 Ts to strands a and a*, 6 Ts to strands b and b*, 8 Ts to strands c and c* and 10 Ts to strands d and d*, where strands a-d correspond to PX and a*-d* correspond to the anti-PX. Reassociation of the PX and anti-PX molecules using the new design will involve the same hybridization partners and base pairs but will however have additional encoding elements (polyTs) that add to the molecular weight. For example, reassociation of the component strands of PX and anti-PX involves the same number of base pairs that hybridize (38 bp), but each of the duplexes will have different number of polyTs that define the encoded bit. If all four strands are encoded, the duplexes that result from reassociation of PX and anti-PX will have molecular weights of 46, 50, 54 and 58 bp respectively due to the additional Ts, providing four bands on a gel corresponding to the 4 bits. Thus, a combination of PX and anti-PX motifs with specific encoding elements allows a hexadecimal readout containing 4 bits (**Fig. 1e**). Strands that do not contain encoding elements reassociate into 38 bp duplexes that are considered to be in a “bin” and do not contribute to encoding (**Fig. 1f**).

First, we confirmed proper assembly of the PX and anti-PX structures containing all four encoding elements. Non-denaturing gels showed the formation of a single dominant band that corresponds to the target structure. The PX and anti-PX with the encoding elements migrated slower compared to the uncoded PX and anti-PX, consistent with the additional nucleotides in the structures (**Fig 2a** and **Fig. S2**). Since our readout is based on gel electrophoresis, one of the criteria is the resolution of the gel to distinctly separate the four bits. To achieve this, we tested the separation of the four different duplex bits on gels containing different percentages of polyacrylamide (**Fig 2b**). We analyzed this result by plotting the migration of the complexes as a Ferguson plot and chose the 14% for further experiments to resolve the four bits (**Fig 2c-d** and **Fig. S3-S4**). While the same slopes are expected for duplexes in such a plot, we observed slight differences that we attribute to the single stranded polyT overhangs on the duplexes.

**Figure 2.**
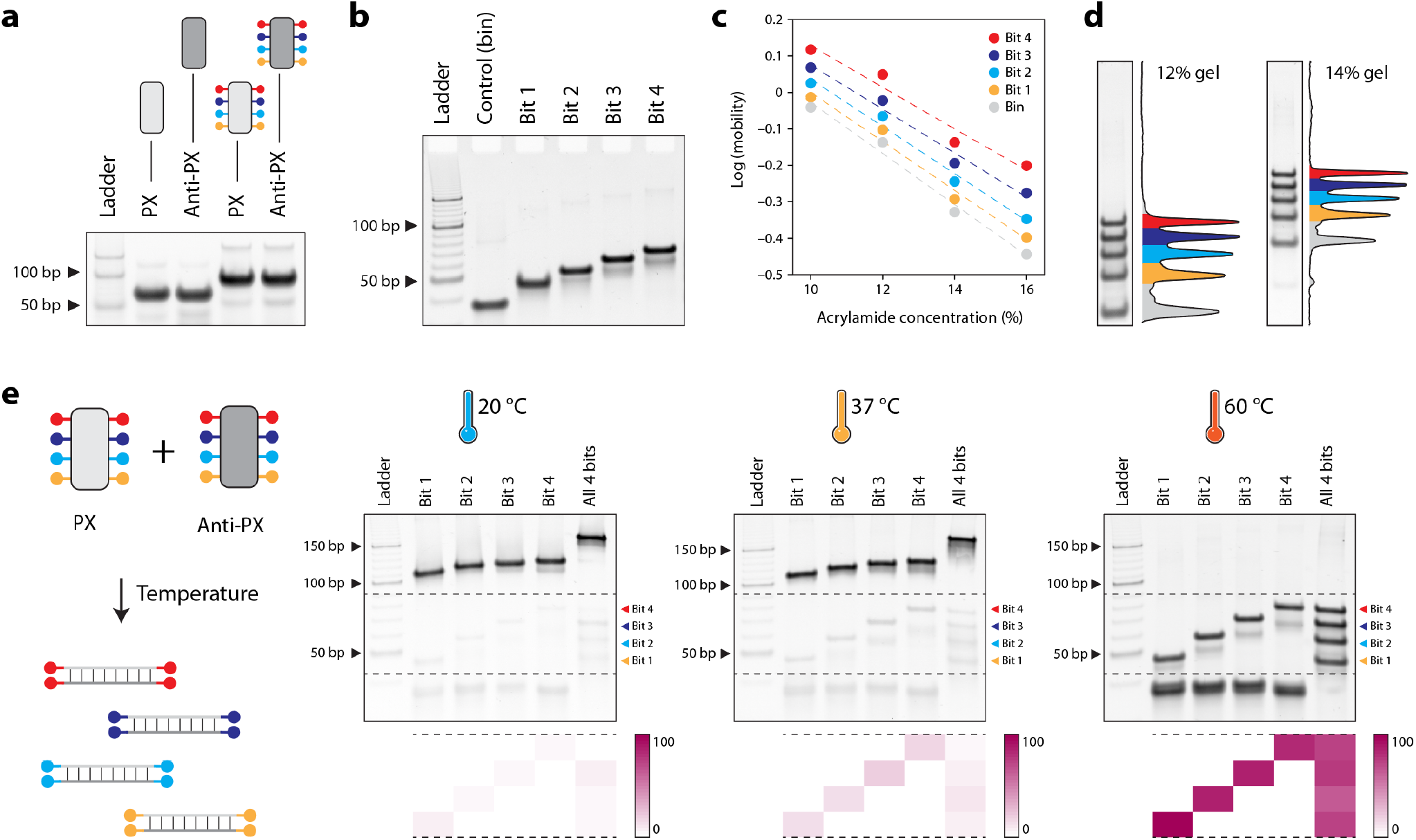
PX DNA assembly and encryption. (a) Assembly of PX and anti-PX motifs with and without encoding elements. (b) Migration analysis of duplex products corresponding to the four bits. (c) Analysis of bit migration across different gel percentages. (d) Visualization of all four bits in a single lane and corresponding band intensities. (e) Decryption by reassociation of PX DNA into duplex bits at different temperatures.

Next, we assembled PX and anti-PX structures that will result in one of the 4 bits or all four. We tested reassociation of these complexes at 20 °C, 37 °C and 60 °C, expecting proper reassociation only at 60 °C. We tested the reassociated samples on a non-denaturing gel and observed the appearance of the four bands corresponding to the 4 bits only when reassociation was performed at 60 °C, although a slight portion of some of the bits was observed at the lower temperatures (**Fig 2e** and **Fig. S5**). Quantified results showed that each of the 4 bits were present as the dominant signal only when one of them was the intended result. In the sample containing all four encoding elements, the resulting signals were similar across the different bits.

Stability of DNA nanostructures is a key parameter in data storage and encryption.^40^ To evaluate the shelf life of the stored information, we mixed the data-encoded PX and anti-PX corresponding to one each of the four bits and stored them for ~4 days at 20 °C, 37 °C or outdoors (temperature range from 20 to 30 °C, **Table S2**). We observed that the information remained almost fully encrypted, as the PX and anti-PX structures do not reassociate at these storage temperatures (**Fig. 3a** and **Fig. S6**). However, when the same mixture was treated at 60 °C, we obtained the encoded four bits. These results demonstrate that we can encode information as four bits in PX DNA structure that is stable over several days and decrypted only with a key (heat to 60 °C). An additional feature of PX DNA is its enhanced nuclease resistance,^41^ providing a higher biostability for the encoded information compared to the decrypted information. We show that the data-encoded PX/anti-PX mixture has an enhanced nuclease resistance when tested against the common nuclease DNase I compared to the decrypted four bits (**Fig. 3b** and **Fig. S7**).

**Figure 3.**
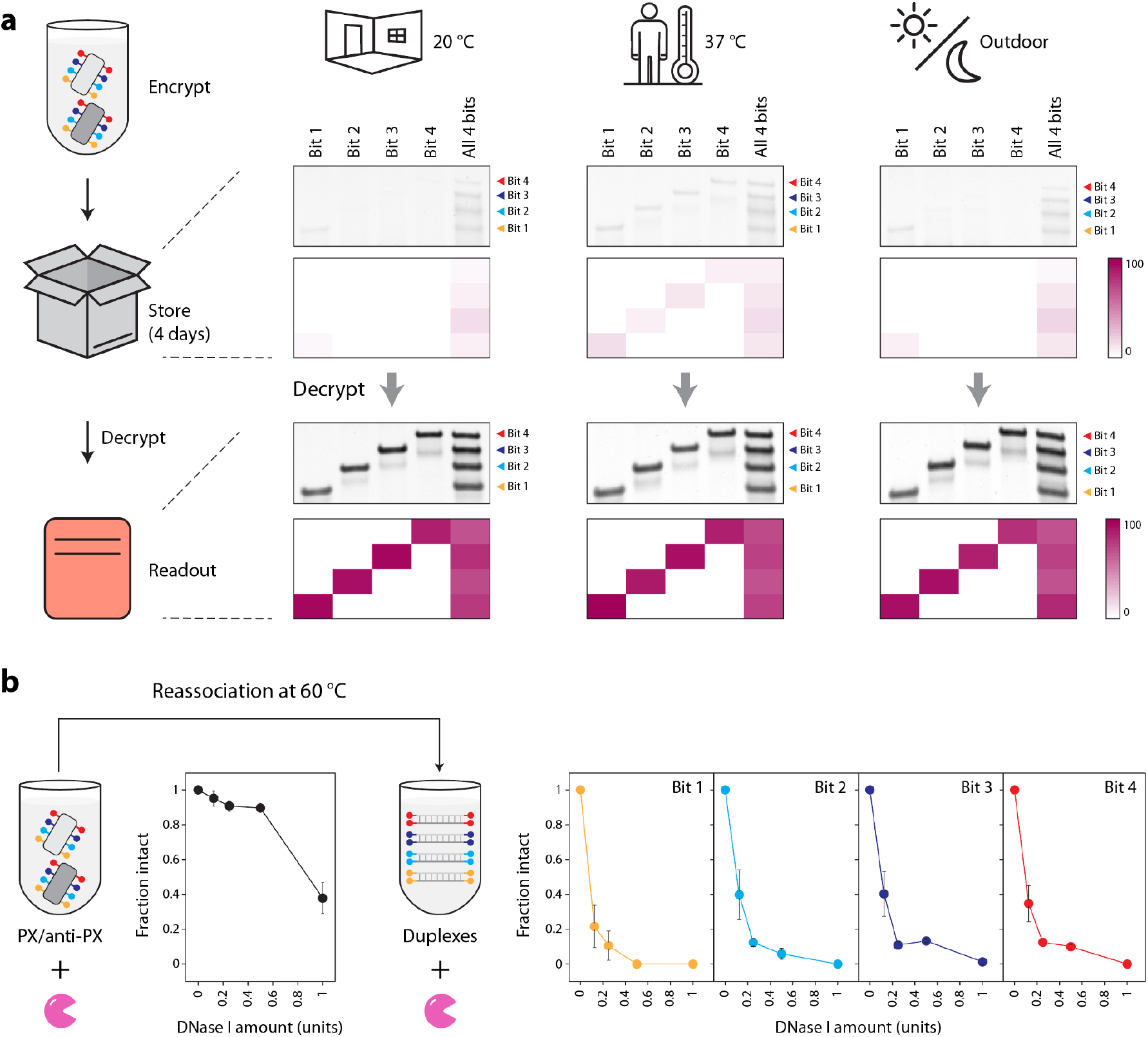
Shelf life and biostability of encoded information. (a) Data-encoded PX and anti-PX DNA stored at different conditions for ~4 days and decrypted at 60 °C. Encoded bits are properly readout only after the heat treatment. (b) Nuclease resistance analysis of encoded PX/anti-PX mixture and the resulting duplexes.

Next, we pursued the encryption and decryption of information represented in hexadecimals. The four strands of the PX DNA structure can each be modified to contain an encoding element, allowing 4 encoding elements (or bits) in a given structure. This design provides a total of 16 combinations of the encoding elements that we encrypt in hexadecimals. We assembled the PX and anti-PX structures corresponding to each of the 16 combinations and validated assembly using non-denaturing PAGE (**Fig 4a, Fig. S8** and **Table S3**). When combined, the PX and anti-PX structures contain encrypted data that is decrypted by treating them at 60 °C, followed by a gel readout of hexadecimal data including the numerals 0-9 and alphabet letters A-F (**Fig 4b** and **Fig. S9**). As a reminder, in cases where fewer than 4 bits are encoded, the PX and anti-PX strands that do not have any encoding elements form duplexes that are considered to be in the “bin”, not contributing to the four bits (two examples shown in **Fig. 4b**). We use this encryption system to demonstrate encoding of words (eg: SCAFFOLD and DATABASE) and hexadecimal codes of colors (eg: dark cyan, #1396AD and soft green, #B7EB65) (**Fig 4c** and **Fig. S10-11**).

**Figure 4.**
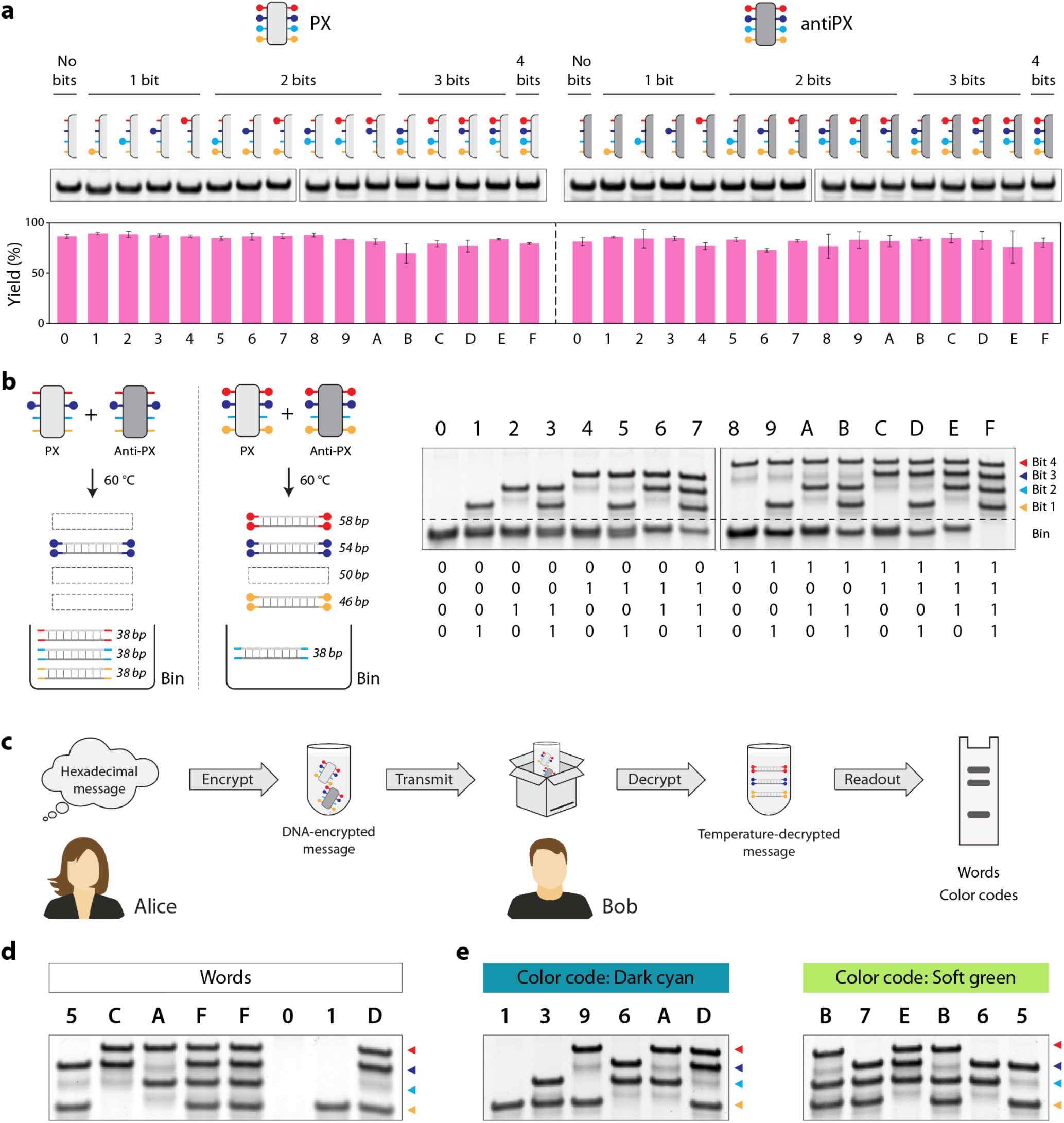
Hexadecimal encryption in PX DNA. (a) Assembly of PX and anti-PX DNA with all combinations of encoding elements. (b) Hexadecimal codes and read out. (c) Strategy to encrypt information represented in hexadecimal codes and decryption at a specific temperature followed by a gel read out of words (d) and color codes (e).

Overall, we show data encoding within DNA nanostructures using PX DNA as a model system. Our strategy allows information encoding as binary bits with up to 4 bits per structure, allowing 16 combinations that can be used as a hexadecimal data system. The encoded structures are stable over several days when stored and the encrypted information is readout only after decryption at 60 °C. Some limitations of our work include a low level of PX/anti-PX reassociation of the bits at temperatures lower than the decryption temperature, requirement of a library of structures with different encoding elements and a 4-bit limitation per lane. The leakage in reassociation can be potentially addressed by careful sequence design for the PX structures, used of other motifs since reassociation is melting temperature-dependent,^39^ and the use of different encoding sequences instead of polyTs. A combination of structure-based reassociation and the toehold-based initiation might also be useful in this regard. While our encoding method requires libraries of PX and anti-PX structures, this is not a limitation unique to our system as almost all DNA-based data storage systems require either a library of specific DNA sequences or structures.

Our work can also be modified to include DNA or RNA encoding elements that are processed by target-specific enzymes to provide additional control on bit readout and erasure^42–44^ and combined with prior strategies that use specific experimental conditions such as DNA concentrations and temperatures to select a subset of structures for encryption from a pool of encoded structures.^45^ While temperature control has been a key feature in programmable DNA synthesis from structured building blocks^46^ and for compartmentalization of reactions in DNA data storage and retrieval,^47^ temperature is rarely used with structural encoding due to the denaturing of assembled structures at high temperatures. However, our strategy uses temperature-dependent bit decryption and might thus be compatible with the other temperature-based processes. In prior work, single stranded extensions on ss-dsDNA encoding bits were used as handles to separate molecules involved in the same file,^48^ a strategy that can also be potentially implemented here. Our strategy can be expanded to 6 bits using the triple-helical domain PX DNA that contains six strands, allowing a total combination of 64 characters. Functionalization of the component strands with fluorophores would also allow expansion of the number of bits and multiplexed readout. Encoding information in such DNA structures expands the use of DNA nanotechnology in molecular computation and materials science.

## Supporting information

Experimental procedures, additional results and DNA sequences used.

## Author contributions

A.K. designed experiments, performed experiments and analyzed data. A.R.C. conceptualized and supervised the project, designed experiments, analyzed data, visualized data and wrote the paper.

## Acknowledgments

Research reported in this publication was supported by the National Institutes of Health (NIH) through National Institute of General Medical Sciences (NIGMS) under award number R35GM150672 to A.R.C. The NSF REU program at The RNA Institute, University at Albany provided summer support for A.K.

## References

(1) Zhirnov, V.; Zadegan, R. M.; Sandhu, G. S.; Church, G. M.; Hughes, W. L. Nucleic Acid Memory.Nature Materials 2016, 15 (4), 366–370. 10.1038/nmat4594.

(2) Dickinson, G. D.; Mortuza, G. M.; Clay, W.; Piantanida, L.; Green, C. M.; Watson, C.; Hayden, E. J.; Andersen, T.; Kuang, W.; Graugnard, E.; Zadegan, R.; Hughes, W. L. An Alternative Approach to Nucleic Acid Memory. Nat Commun 2021, 12 (1), 2371. 10.1038/s41467-021-22277-y.

(3) Zhang, Y.; Ren, Y.; Liu, Y.; Wang, F.; Zhang, H.; Liu, K. Preservation and Encryption in DNA Digital Data Storage. ChemPlusChem 2022, 87 (9), e202200183. 10.1002/cplu.202200183.

(4) Ceze, L.; Nivala, J.; Strauss, K. Molecular Digital Data Storage Using DNA. Nat Rev Genet 2019, 20(8), 456–466. 10.1038/s41576-019-0125-3.

(5) Wang, J.; Xie, M.; Ouyang, L.; Li, J.; Wang, L.; Fan, C.; Chao, J. Artificial Molecular Communication Network Based on DNA Nanostructures Recognition. Nat Commun 2025, 16 (1), 244. 10.1038/s41467-024-55527-w.

(6) Erlich, Y.; Zielinski, D. DNA Fountain Enables a Robust and Efficient Storage Architecture. Science 2017, 355 (6328), 950–954. 10.1126/science.aaj2038.

(7) Church, G. M.; Gao, Y.; Kosuri, S. Next-Generation Digital Information Storage in DNA. Science 2012, 337 (6102), 1628–1628. 10.1126/science.1226355.

(8) Antkowiak, P. L.; Lietard, J.; Darestani, M. Z.; Somoza, M. M.; Stark, W. J.; Heckel, R.; Grass, R. N. Low Cost DNA Data Storage Using Photolithographic Synthesis and Advanced Information Reconstruction and Error Correction. Nat Commun 2020, 11 (1), 5345. 10.1038/s41467-020-19148-3.

(9) Chen, K.; Kong, J.; Zhu, J.; Ermann, N.; Predki, P.; Keyser, U. F. Digital Data Storage Using DNA Nanostructures and Solid-State Nanopores. Nano Lett. 2019, 19 (2), 1210–1215. 10.1021/acs.nanolett.8b04715.

(10) Song, L.; Wang, G.; Wei, Y.; Huang, Y.; Zhang, Y.; Liu, X.; Yuchi, Z.; Yuan, Y.; Luo, Y.; Zhang, Y. ZAT-DNA Enables DNA Data Storage with Molecular-Layer Non-Replicability. Nat Commun 2026. 10.1038/s41467-026-72869-9.

(11) Fan, C.; Deng, Q.; Zhu, T. F. Bioorthogonal Information Storage in L-DNA with a High-Fidelity Mirror-Image Pfu DNA Polymerase. Nat Biotechnol 2021, 39 (12), 1548–1555. 10.1038/s41587-021-00969-6.

(12) Zhang, C.; Wu, R.; Sun, F.; Lin, Y.; Liang, Y.; Teng, J.; Liu, N.; Ouyang, Q.; Qian, L.; Yan, H. Parallel Molecular Data Storage by Printing Epigenetic Bits on DNA. Nature 2024, 634 (8035), 824–832. 10.1038/s41586-024-08040-5.

(13) Fan, Q.; Zhao, X.; Li, J.; Liu, R.; Liu, M.; Feng, Q.; Long, Y.; Fu, Y.; Zhai, J.; Pan, Q.; Li, Y. De Novo Non-Canonical Nanopore Basecalling Enables Private Communication Using Heavily-Modified DNA Data at Single-Molecule Level. Nat Commun 2025, 16 (1), 4099. 10.1038/s41467-025-59357-2.

(14) Tabatabaei Yazdi, S. M. H.; Yuan, Y.; Ma, J.; Zhao, H.; Milenkovic, O. A Rewritable, Random-Access DNA-Based Storage System. Sci Rep 2015, 5 (1), 14138. 10.1038/srep14138.

(15) Wang, J.; Gao, R.; Zhang, J.; Wang, H.; Luo, Z.; Fan, C.; Chao, J.; Wang, L. A Multiple-Encrypted DNA Device for Secure Communication. Science Advances 2026, 12 (27), eaef9101. 10.1126/sciadv.aef9101.

(16) Halvorsen, K.; Wong, W. P. Binary DNA Nanostructures for Data Encryption. PLOS ONE 2012, 7 (9), e44212. 10.1371/journal.pone.0044212.

(17) Takinoue, M.; Suyama, A. Hairpin-DNA Memory Using Molecular Addressing. Small 2006, 2 (11), 1244–1247. 10.1002/smll.200600237.

(18) Garibotti, A. V.; Liao, S.; Seeman, N. C. A Simple DNA-Based Translation System. Nano Lett. 2007,7 (2), 480–483. 10.1021/nl0628605.

(19) Zhang, C.; Xie, M.; Wang, L.; Fan, C.; Chao, J. Linked Data Storage Using DNA Origami Nanostructures. Nat Commun 2025, 16 (1), 11422. 10.1038/s41467-025-66274-x.

(20) Chu, L.; Wan, L.; Wang, H.; Wang, P.; Deng, L.; Gao, Y.; Guo, P.; He, L.; Han, D. Hierarchical Core– Shell DNA Condensates Enable Programmable Information Storage and Encryption. Nat Commun 2025, 17 (1), 401. 10.1038/s41467-025-67093-w.

(21) Shin, J.-S.; Pierce, N. A. Rewritable Memory by Controllable Nanopatterning of DNA. Nano Lett.2004, 4 (5), 905–909. 10.1021/nl049658r.

(22) Chandrasekaran, A. R.; Trivedi, R.; Halvorsen, K. Ribonuclease-Responsive DNA Nanoswitches. Cell Reports Physical Science 2020, 1 (7), 100117. 10.1016/j.xcrp.2020.100117.

(23) Chen, K.; Fan, S.; Liu, N.; Song, J.; Pan, L. Hierarchical Access to Encoded Data on DNA Nanostructures Using Administrator and User Keys. Nucleic Acids Res 2025, 53 (16), gkaf835. 10.1093/nar/gkaf835.

(24) Chandrasekaran, A. R.; Levchenko, O.; Patel, D. S.; MacIsaac, M.; Halvorsen, K. Addressable Configurations of DNA Nanostructures for Rewritable Memory. Nucleic Acids Res 2017, 45 (19), 11459–11465. 10.1093/nar/gkx777.

(25) Chandrasekaran, A. R. Reconfigurable DNA Nanoswitches for Graphical Readout of Molecular Signals. ChemBioChem 2018, 19 (10), 1018–1021. 10.1002/cbic.201800057.

(26) Talbot, H.; Halvorsen, K.; Chandrasekaran, A. R. Encoding, Decoding, and Rendering Information in DNA Nanoswitch Libraries. ACS Synth. Biol. 2023, 12 (4), 978–983. 10.1021/acssynbio.2c00649.

(27) Forrest, N. T.; Vilcapoma, J.; Alejos, K.; Halvorsen, K.; Chandrasekaran, A. R. Orthogonal Control of DNA Nanoswitches with Mixed Physical and Biochemical Cues. Biochemistry 2021, 60 (4), 250–253. 10.1021/acs.biochem.0c00952.

(28) Chandrasekaran, A. R.; Punnoose, J. A.; Valsangkar, V.; Sheng, J.; Halvorsen, K. Integration of a Photocleavable Element into DNA Nanoswitches. Chem. Commun. 2019, 55 (46), 6587–6590. 10.1039/C9CC03069G.

(29) Wisna, G. B. M.; Sukhareva, D.; Zhao, J.; Chopade, P.; Satyabola, D.; Matthies, M.; Roy, S.; Wang, C.; Šulc, P.; Yan, H.; Hariadi, R. F. High-Speed 3D DNA PAINT and Unsupervised Clustering for Unlocking 3D DNA Origami Cryptography. Nat Commun 2025, 16 (1), 11514. 10.1038/s41467-025-66338-y.

(30) Shih, W. M.; Quispe, J. D.; Joyce, G. F. A 1.7-Kilobase Single-Stranded DNA That Folds into a Nanoscale Octahedron. Nature 2004, 427 (6975), 618. 10.1038/nature02307.

(31) Ohayon, Y. P.; Sha, R.; Flint, O.; Liu, W.; Chakraborty, B.; Subramanian, H. K. K.; Zheng, J.;Chandrasekaran, A. R.; Abdallah, H. O.; Wang, X.; Zhang, X.; Seeman, N. C. Covalent Linkage of One-Dimensional DNA Arrays Bonded by Paranemic Cohesion. ACS Nano 2015, 9 (10), 10304–10312. 10.1021/acsnano.5b04335.

(32) Shen, W.; Liu, Q.; Ding, B.; Shen, Z.; Zhu, C.; Mao, C. The Study of the Paranemic Crossover (PX) Motif in the Context of Self-Assembly of DNA 2D Crystals. Org. Biomol. Chem. 2016, 14 (30), 7187–7190. 10.1039/C6OB01146B.

(33) Yan, H.; Zhang, X.; Shen, Z.; Seeman, N. C. A Robust DNA Mechanical Device Controlled by Hybridization Topology. Nature 2002, 415 (6867), 62. 10.1038/415062a.

(34) Gu, H.; Chao, J.; Xiao, S.-J.; Seeman, N. C. A Proximity-Based Programmable DNA Nanoscale Assembly Line. Nature 2010, 465 (7295), 202–205. 10.1038/nature09026.

(35) Zhong, H.; Seeman, N. C. RNA Used to Control a DNA Rotary Nanomachine. Nano Lett. 2006, 6 (12), 2899–2903. 10.1021/nl062183e.

(36) Chakraborty, B.; Sha, R.; Seeman, N. C. A DNA-Based Nanomechanical Device with Three Robust States. PNAS 2008, 105 (45), 17245–17249. 10.1073/pnas.0707681105.

(37) Chandrasekaran, A. R. A DNA Rotary Nanodevice Operated by Enzyme-Initiated Strand Resetting. Chem. Commun. 2024, 60 (5), 534–537. 10.1039/D3CC05487J.

(38) Talbot, H.; Chandrasekaran, A. R. Mismatch-Induced Toehold-Free Strand Displacement Used to Control a DNA Nanodevice. ACS Synth. Biol. 2025, 14 (6), 1931–1935. 10.1021/acssynbio.5c00107.

(39) Kabbara, N.; Anderson, L. A.; Singha, S.; Chandrasekaran, A. R. Controlled Reassociation of Multistranded, Polycrossover DNA Molecules into Double Helices. Nano Lett. 2025, 25 (47), 16658–16663. 10.1021/acs.nanolett.5c04286.

(40) Matange, K.; Tuck, J. M.; Keung, A. J. DNA Stability: A Central Design Consideration for DNA Data Storage Systems. Nat Commun 2021, 12 (1), 1358. 10.1038/s41467-021-21587-5.

(41) Chandrasekaran, A. R.; Vilcapoma, J.; Dey, P.; Wong-Deyrup, S. W.; Dey, B. K.; Halvorsen, K. Exceptional Nuclease Resistance of Paranemic Crossover (PX) DNA and Crossover-Dependent Biostability of DNA Motifs. J. Am. Chem. Soc. 2020, 142 (14), 6814–6821. 10.1021/jacs.0c02211.

(42) Zhang, J.; Hou, C.; Liu, C. CRISPR-Powered Quantitative Keyword Search Engine in DNA Data Storage. Nat Commun 2024, 15 (1), 2376. 10.1038/s41467-024-46767-x.

(43) Faheem, H.; Mathivanan, J.; Talbot, H.; Zeghal, H.; Vangaveti, S.; Sheng, J.; Chen, A. A.; Chandrasekaran, A. R. Toehold Clipping: A Mechanism for Remote Control of DNA Strand Displacement. Nucleic Acids Research 2023, 51 (8), 4055–4063. 10.1093/nar/gkac1152.

(44) Imburgia, C.; Organick, L.; Zhang, K.; Cardozo, N.; McBride, J.; Bee, C.; Wilde, D.; Roote, G.; Jorgensen, S.; Ward, D.; Anderson, C.; Strauss, K.; Ceze, L.; Nivala, J. Random Access and SemanticSearch in DNA Data Storage Enabled by Cas9 and Machine-Guided Design. Nat Commun 2025, 16 (1), 6388. 10.1038/s41467-025-61264-5.

(45) Tomek, K. J.; Volkel, K.; Indermaur, E. W.; Tuck, J. M.; Keung, A. J. Promiscuous Molecules for Smarter File Operations in DNA-Based Data Storage. Nat Commun 2021, 12 (1), 3518. 10.1038/s41467-021-23669-w.

(46) Kim, J.; Choi, J.; Kim, W.; Cho, N.; Jung, Y.; Park, S.; Choi, E.; Lee, C.; Kim, Y.; Ryu, T.; Choi, H.; Choi, Y. Programmable One-Pot Polymerase-Mediated DNA Synthesis via Temperature Control. Nat Commun 2026. 10.1038/s41467-026-74890-4.

(47) Bögels, B. W. A.; Nguyen, B. H.; Ward, D.; Gascoigne, L.; Schrijver, D. P.; Makri Pistikou, A.-M.; Joesaar, A.; Yang, S.; Voets, I. K.; Mulder, W. J. M.; Phillips, A.; Mann, S.; Seelig, G.; Strauss, K.; Chen, Y.-J.; de Greef, T. F. A. DNA Storage in Thermoresponsive Microcapsules for Repeated Random Multiplexed Data Access. Nat. Nanotechnol. 2023, 18 (8), 912–921. 10.1038/s41565-023-01377-4.

(48) Lin, K. N.; Volkel, K.; Tuck, J. M.; Keung, A. J. Dynamic and Scalable DNA-Based Information Storage. Nat Commun 2020, 11 (1), 2981. 10.1038/s41467-020-16797-2.

