## Supporting information for "Hexadecimal data encryption in paranemic crossover (PX) DNA"

### MATERIALS AND METHODS

#### Preparation of DNA complexes

DNA strands were purchased from Integrated DNA Technologies (IDT). Full sequences are listed in Table S1. PX DNA structures were prepared by mixing the component DNA strands in equal ratios in Tris-Acetic-EDTA buffer containing 40 mM Tris base (pH 8), 20 mM acetic acid, 2 mM EDTA, and 12.5 mM magnesium acetate (1× TAE-$Mg^{2+}$). Samples were annealed from 90 °C to 20 °C over 48 h. For duplexes, strands were combined in equal ratios in 1× TAE-$Mg^{2+}$ buffer and annealed from 90 °C to 20 °C over 30 minutes. For structure reassociation, pre-assembled PX and anti-PX structures were mixed in equal molar ratios and incubated at different temperatures in a BioRad thermal cycler or on the bench (20 °C) for 3 h.

#### DNA strand combination for encoding bits

Bits are encoded in any of the four strands of the PX and aniti-PX. PX and anti-PX structures with the same combination of encoding elements are used for encryption, resulting in distinct duplex bits. For bit 1, strand a contains the polyTs and strands b, c, d do not contain any poly Ts. Similarly, strands b, c and d containing polyTs are used when encoding bits 2, 3 and 4 respectively. Any combination of these four strands are used for the hexadecimal representation. Strands that do not contain the encoding elements form a 38 bp duplex that does not encode any bits and is considered to be in the "bin" (see Figure 1f).

#### Temperature-based decryption

Encoded PX and anti-PX structures were mixed in equimolar ratios and incubated at 20 °C, 37 °C or 60 °C for 3 h. The decryption temperature was chosen to be 60 °C, with a typical incubation time of 3 h. Once decrypted, the samples remain decrypted (i.e. in the duplex form) and can be read out using gel electrophoresis. For shelf life experiments, PX and anti-PX structures with specific combinations of encoding elements (bits 1, 2, 3, 4 and all four bits together) were mixed and incubated at 20 °C (on the bench), 37 °C (in a thermal cycler) or outdoors in a tube rack. Outdoor temperatures were monitored and are provided in Table S2.

#### Nuclease degradation assay

For PX samples, the PX and anti-PX were mixed at a final concentration of 0.5 μM and used for the DNase I assay (DNase I, New England Biolabs). To convert the PX/anti-PX mixture into duplexes, the PX and anti-PX mixture was heated at 60 °C for 3 h and the resulting sample was used in the DNase I assay. DNA samples were first mixed with DNase I reaction buffer (provided by the vendor) to a final 1× concentration. Enzyme dilutions were made in nuclease-free water. For the nuclease degradation assay, 1 μl of the enzyme was added to 8 μl of the DNA sample containing the reaction buffer and incubated at 20 °C for 30 min. Incubated samples were mixed with gel loading dye containing bromophenol blue and 1× TAE-$Mg^{2+}$ buffer and run on non-denaturing gels to analyze degradation. Degradation profiles were obtained by normalizing the band intensity corresponding to the structure (at each enzyme concentration) to the control lane without any enzyme. For the encrypted (PX/anti-PX) sample, the intensity was normalized to the band in the lane without any nuclease. For the four-bit read out, the bands corresponding to each of the four bits was normalized to the starting band intensity (without enzyme) of that particular bit.

**Hexadecimal representation**

Words or color codes were chosen, and the corresponding 4-bit binary code was used to assemble the PX and anti-PX structures containing the encoding elements. Binary codes for hexadecimal representation are provided in Table S3. PX and anti-PX mixtures are considered encrypted, and the information is decrypted by heating the samples at 60 °C for 3 h, followed by a gel read out (eg: Figure 4d-4e).

**Gel electrophoresis and quantification**

Non-denaturing gels were prepared using 19:1 acrylamide/bisacrylamide (National Diagnostics). Samples were mixed with loading dye containing bromophenol blue and glycerol prior to loading. Gels were run at a constant voltage at 4 °C in 1× TAE-$Mg^{2+}$ running buffer. Gels were stained in 0.5× GelRed (Biotium), imaged using a Bio-Rad Gel Doc XR+ and analyzed using ImageLab or Image J software. Images were typically taken at multiple exposures ranging from 3 to 30 s to facilitate accurate quantification. For each gel, quantification was done using the highest-exposure image that did not contain saturated pixels in the bands. For analysis of replicates, separate gels with the same exposure time were typically used for quantifying band brightness. The assembly yield was quantified as the fraction of the intensity corresponding to the band of interest compared to the total intensity of all the bands in the lane. For the heat maps in Figures 2e and 3a, the measured intensities were normalized to the highest band intensity of the four bits in the specific experiment.

**Gel migration analysis**

Migration of the bands were measured using Image J software as distance travelled by the band in the gel with the well as the reference point. 10, 12, 14 and 16% polyacrylamide gel percentages were used. Gels were imaged at the same dimensions to allow migration analysis across gels. Migration distances were plotted as log (mobility) vs gel % to obtain a Ferguson plot to compare the four bits on a gel read out (Figure 2b-2c and Figure S3). Area under the curve when obtaining band intensities are used to show the migration of the four bits in Figure 2d using two representative gels: 12% and 14% (Figure S4).

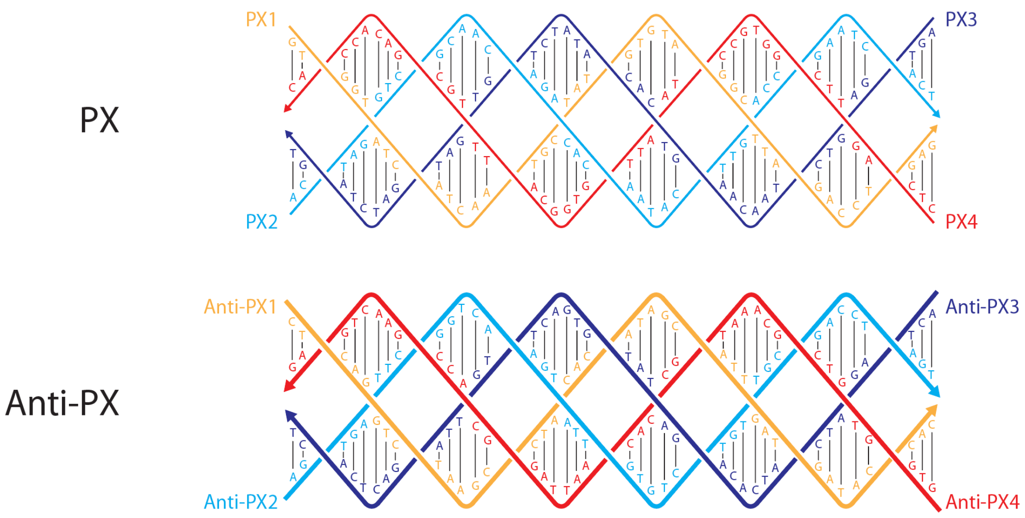


**Figure S1**. Sequences of the PX and anti-PX motifs.

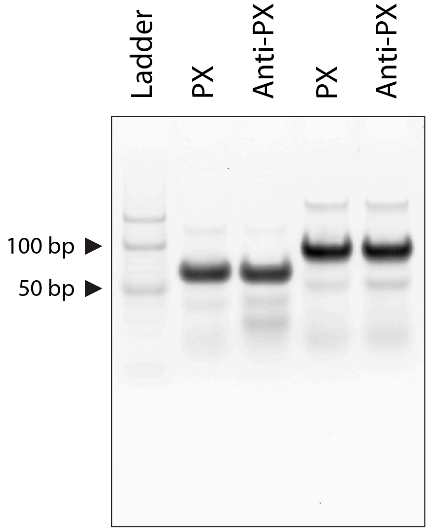


**Figure S2**. Non-denaturing gel showing the assembly of PX and anti-PX motifs with and without all four encoding elements present. Full image of gel shown in Figure 2a.

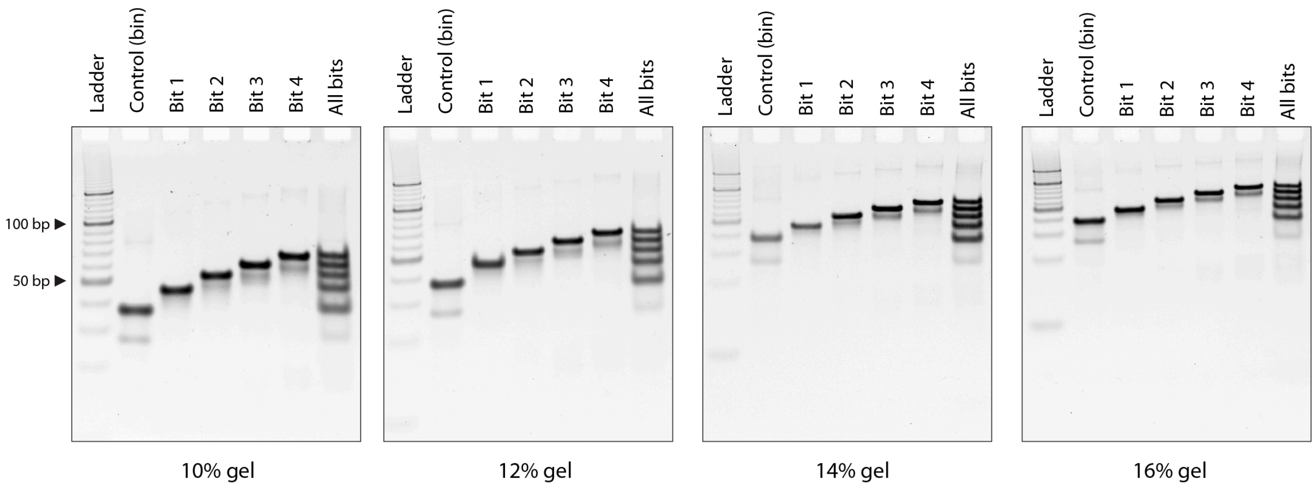


**Figure S3**. Migration of different bits in different polyacrylamide percentages. Full image of gel shown in Figure 2b and gels used for plot shown in Figure 2c.

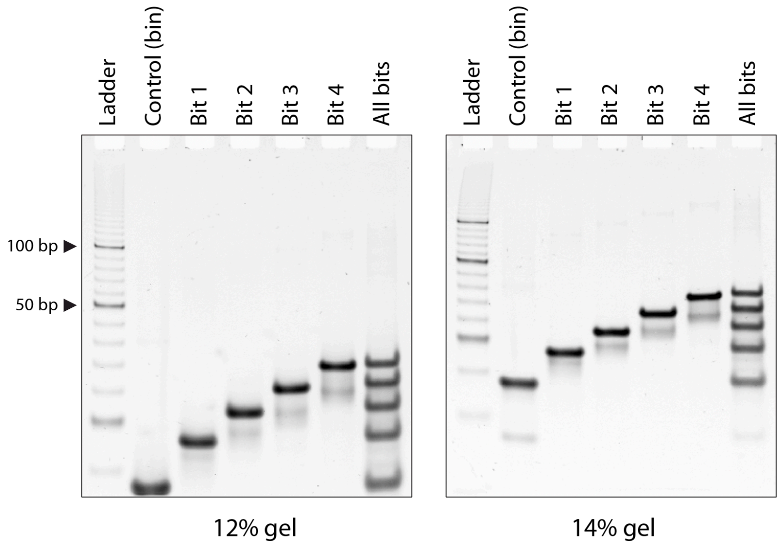


**Figure S4**. Additional optimization of gel percentage and run time. Full images of gels shown in Figure 2d.

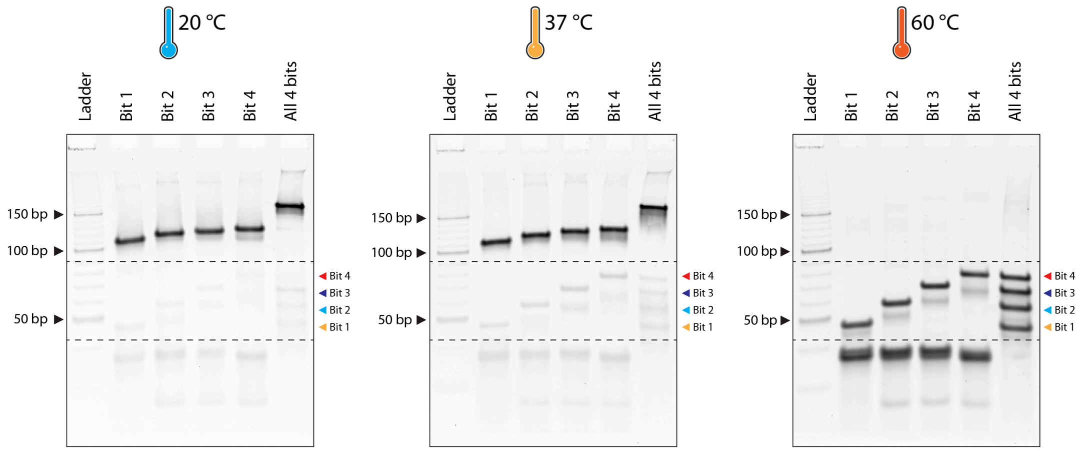


**Figure S5**. Temperature-dependent reassociation of PX structures into duplexes for different bits. Full images of gels shown in Figure 2e.

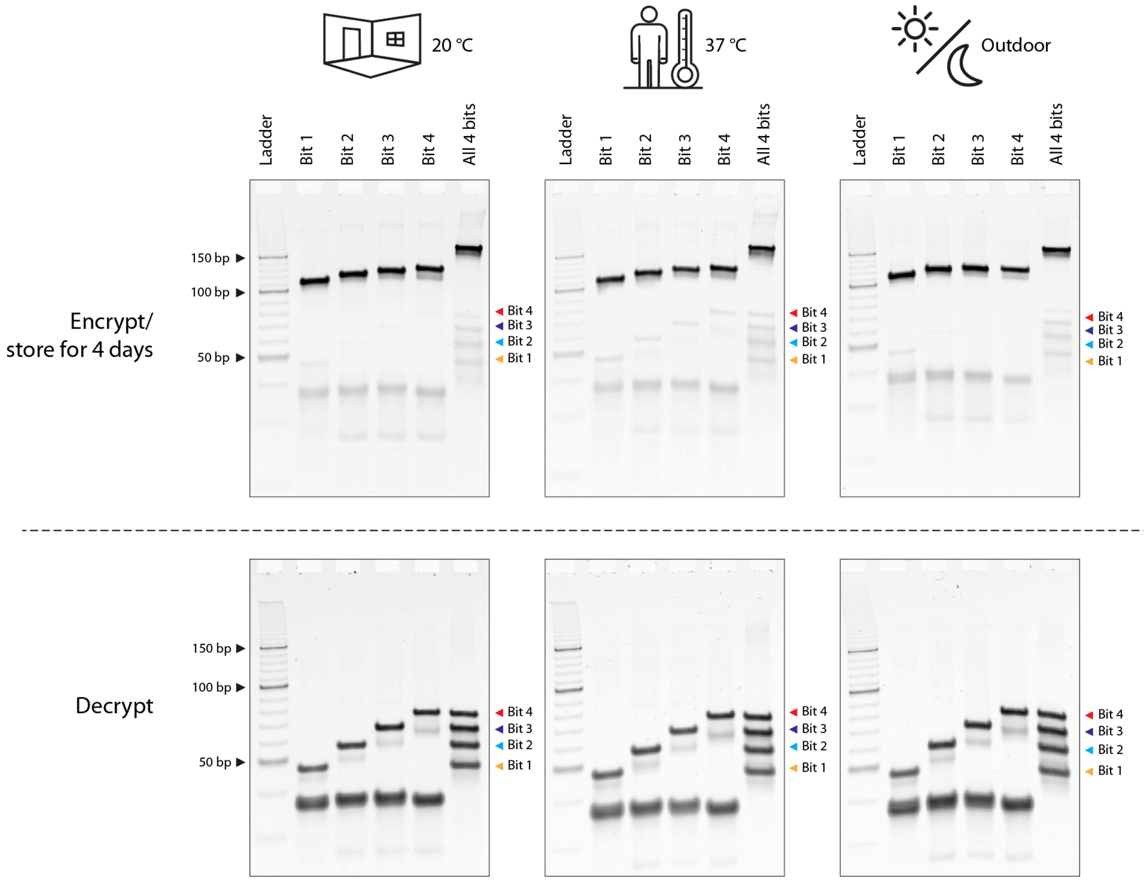


**Figure S6**. Shelf life of PX-encrypted data. Top row shows DNA samples stored in different conditions for ~4 days. Bottom row shows the same stored samples after treatment at 60 °C for 3 h. Full images of gels shown in Figure 3a.

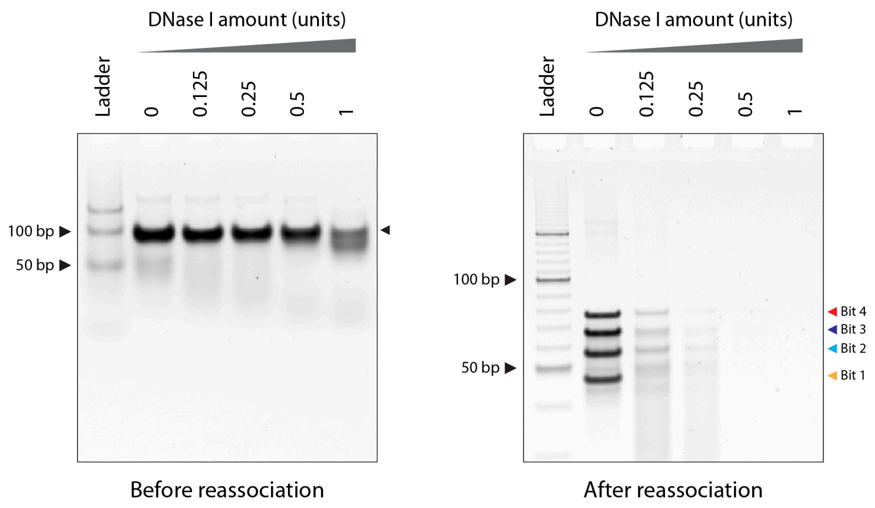


**Figure S7**. Nuclease resistance of data. Gels showing DNase I treated samples and their degradation in different nuclease amounts. Gel images used for data shown in Figure 3b.

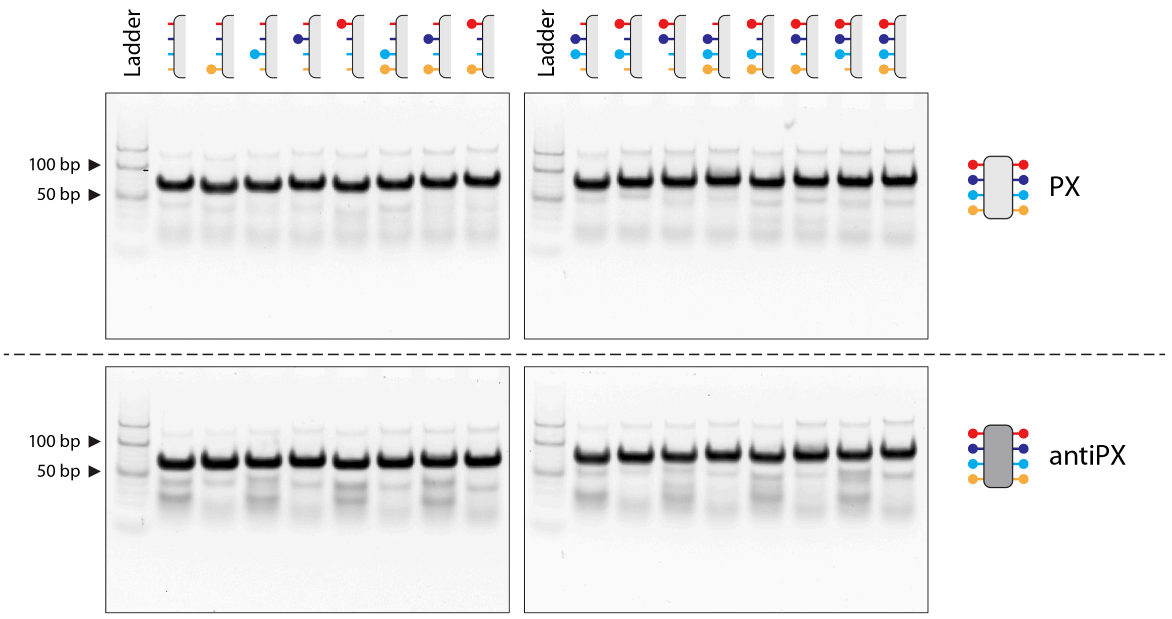


**Figure S8**. Gels showing assembly of all the combinations of PX and anti-PX motifs with encoding elements. Full images of gels shown in Figure 4a.

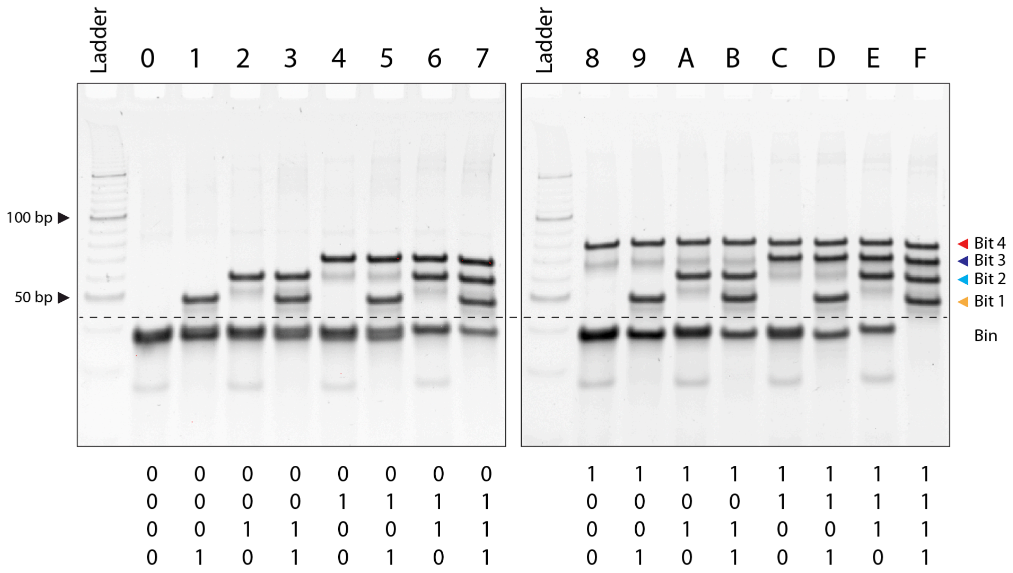


**Figure S9**. Assembly of hexadecimal characters. Full image of gel shown in Figure 4b.

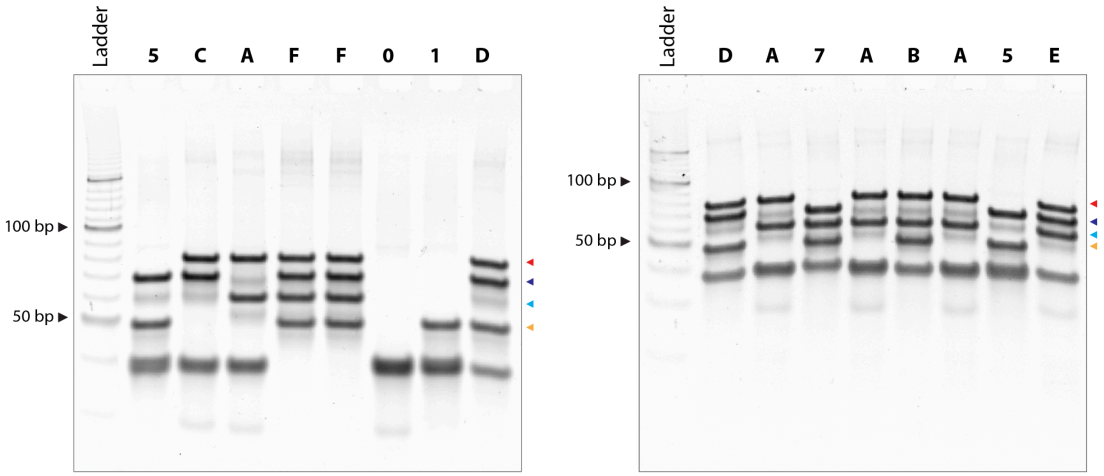


**Figure S10**. Assembly of words in hexadecimal. Full image of gel shown in Figure 4c.

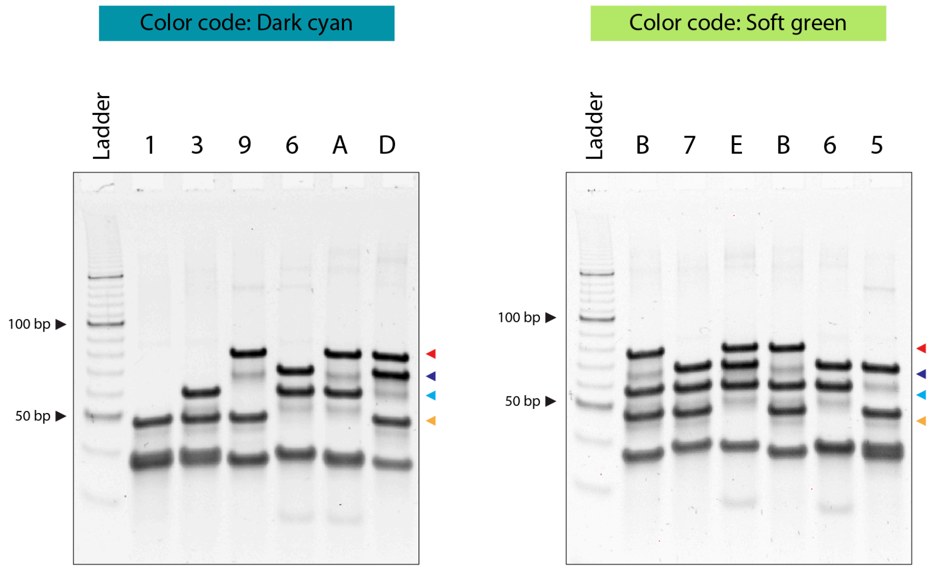


**Figure S11**. Assembly of color codes in hexadecimal. Full image of gel shown in Figure 4c.

| Strand name | Sequence |
|---|---|
| PX-a | GTGGTATCATCAATGCTATGTGTAGGCTTAGACCTGAG |
| PX-b | ACTAGGTCGCAACAGACACAATACTTGACCGAATCACT |
| PX-c | AGTGAGTCTAACAAGTCACATATCTGTGATGATCTAGT |
| PX-d | CTCAGTTCGGTGCCTAATTGTGGCATTTGCGACACCAC |
| Anti-PX-a | CTCAGGTCTAAGCCTACACATAGCATTGATGATACCAC |
| Anti-PX-b | AGTGATTCGGTCAAGTATTGTGTCTGTTGCGACCTAGT |
| Anti-PX-c | ACTAGATCATCACAGATATGTGACTTGTTAGACTCACT |
| Anti-PX-d | GTGGTGTCGCAAATGCCACAATTAGGCACCGAACTGAG |
| PX-a-T4 | TTTTGTGGTATCATCAATGCTATGTGTAGGCTTAGACCTGAGTTTT |
| PX-b-T6 | TTTTTTACTAGGTCGCAACAGACACAATACTTGACCGAATCACTTTTTTT |
| PX-c-T8 | TTTTTTTTAGTGAGTCTAACAAGTCACATATCTGTGATGATCTAGTTTTTTTTT |
| PX-d-T10 | TTTTTTTTTTCTCAGTTCGGTGCCTAATTGTGGCATTTGCGACACCACTTTTTTTTTT |
| Anti-PX-a-T4 | TTTTCTCAGGTCTAAGCCTACACATAGCATTGATGATACCACTTTT |
| Anti-PX-b-T6 | TTTTTTAGTGATTCGGTCAAGTATTGTGTCTGTTGCGACCTAGTTTTTTT |
| Anti-PX-c-T8 | TTTTTTTTACTAGATCATCACAGATATGTGACTTGTTAGACTCACTTTTTTTTT |
| Anti-PX-d-T10 | TTTTTTTTTTGTGGTGTCGCAAATGCCACAATTAGGCACCGAACTGAGTTTTTTTTTT |

**Table S1.** Sequences used in the study (written 5' to 3').

| | Morning | Noon | Afternoon | Night |
|---|---|---|---|---|
| Day 1 | | | 26.7 | 21.7 |
| Day 2 | 20.6 | 24.4 | 28.3 | 22.2 |
| Day 3 | 21.1 | 26.1 | 28.9 | 22.8 |
| Day 4 | 21.1 | 26.7 | 29.4 | 21.7 |
| Day 5 | 23.3 | | | |

**Table S2.** Range of temperatures during outdoor storage.

| **Hexadecimal** | **Binary** |
|---|---|
| 0 | 0000 |
| 1 | 0001 |
| 2 | 0010 |
| 3 | 0011 |
| 4 | 0100 |
| 5 | 0101 |
| 6 | 0110 |
| 7 | 0111 |
| 8 | 1000 |
| 9 | 1001 |
| A | 1010 |
| B | 1011 |
| C | 1100 |
| D | 1101 |
| E | 1110 |
| F | 1111 |

**Table S3.** Hexadecimal representation using binary codes.
